# Beta and theta oscillations correlate with subjective time during musical improvisation in ecological and controlled settings: a single subject study

**DOI:** 10.1101/2020.11.08.373217

**Authors:** Nicolas Farrugia, Alix Lamouroux, Christophe Rocher, Jules Bouvet, Giulia Lioi

## Abstract

In this paper, we describe the results of a single subject study attempting at a better understanding of the subjective state during musical improvisation. In a first experiment, we setup an ecological paradigm measuring EEG on a musician in free improvised concerts with an audience, followed by retrospective rating of the mental state of the improviser. We introduce Subjective Temporal Resolution (STR), a retrospective rating assessing the instantaneous quantization of subjective timing of the improviser. We identified high and low STR states using Hidden Markov Models in two performances, and were able to decode those states using supervised learning on instantaneous EEG power spectrum, showing increases in theta and alpha power with high STR values. In a second experiment, we found an increase of theta and beta power when experimentally manipulating STR in a musical improvisation imagery experiment. These results are interpreted with respect to previous research on flow state in creativity, as well as with the temporal processing literature. We suggest that a component of the subjective state of musical improvisation may be reflected in an underlying mechanism related to the subjective quantization of time. We also demonstrate the feasibility of single case studies of musical improvisation using brain activity measurements and retrospective reports, by obtaining consistent results across multiple sessions.

## 1 Introduction

> Improvisation enjoys the curious distinction of being both the most widely practiced of all musical activities, and the least understood and acknowledged. [5]

Fourty years have passed since Derek Bailey wrote these words [5], and musical improvisation has now been widely acknowledged as a model to investigate the neuroscience of creativity [8, 29]. A wealth of studies done the last 15 years have attempted to elucidate the neural correlates of musical improvisation, mostly through hypothesis-driven research, and broadly asking questions of two types: (1) what makes brain activity during improvisation different than other music-related activity, and (2) is there long term plasticity associated with the (expert) practice of improvisation [8]. Much of these hypothesis seem driven by initial accounts proposed by the theoretical framework from Pressing [44, 45], which considers improvisation as a complex activity requiring significant domain-specific expertise related to musical training such as sensorimotor synchronisation, motor planning, procedural memory for accurate sensorimotor execution, as well as a combination of a range of cognitive functions such as long term memory and generative processes involved in creativity [44]. A wealth of neuroscientific studies have confirmed the role of many brain networks such as the executive control network, notably involved in regulating attention and working memory, as well as the default mode network which mediates mental simulation (e.g. mental time travel) and mind wandering [8]. Studies have shown that while the activity in these two networks were traditionally considered as anti-correlated [47], they can operate concurrently during musical improvisation [42]. More recently, authors have proposed that motor and premotor regions are also involved in musical improvisation, possibly managing temporal aspects of performance [7]. Taken together, these studies have brought light on brain areas that are important for musical improvisation, either because they are activated during performance, or because of long term plasticity effects associated with expertise.

Most of the aforementionned studies have used functional MRI in order to shed light on the spatial location of brain networks involved in improvisation. Many other studies have used electroencephalography (EEG) and magnetoencephalography, in order to get a finer temporal understanding of neuronal activity during improvisation. Studies have found improvisation related activity in the alpha (8–12 Hz) and beta (13–30 Hz) frequency range [9, 19, 51] located in prefrontal and medial frontal areas, while other studies have examined brain connectivity [37, 56] or power changes at the sensor level [17, 48, 49].

Another perspective developed in the literature consists in considering musical improvisation as a *subjective state* [8,18,31,54]. Such an angle considers primarily musical improvisation as an instance of flow state. Flow state is defined as “the holistic sensation that people feel when they act with total involvement,” [13], and has been extensively studied with respect to many domains of subjective experience including creativity and aesthetics, but also in other activities requiring full subject engagement such as sports [14]. Flow state has been studied in the context of the musical improvisation (e.g. see [10, 18, 32, 54]. These studies have discussed the involvement of brain networks of spontaneous, endogenous activity such as the default mode network [42], and considers the notion of flow state as central in the phenomenology of musical improvisation [54]. Interview and observations studies have noted that the concepts usually discussed relating musical improvisation and flow state are selflessness, dream-like experiences, modulation in the passage of time, various forms of mental imagery, and a sense of disconnection with reality [6, 54]. Therefore, previous literature suggest that the subjective experience of musical improvisation is rich and complex, as well as idiosyncratic, which motivates the need for qualitative studies using single cases or structured interviews.

Single case studies are quite common in music studies of improvisation, and have attempted at characterizing the creative process [60, 61]. Interview studies using small groups of expert musicians were also done to study group dynamics in musical improvisation [59], as well as subjective assessment of musical qualities [43, 62]. Interestingly, these studies show that musicians do not necessarily agree on what makes a good musical improvisation, which suggest that the study of musical improvisation on single cases might give unique insights on the cognitive and neural basis of creativity. A recent single case study on a internationally acclaimed musical improviser attempted at examining brain activity during improvisation using fMRI using a classical block-design with controlled experimental conditions [6]. Results suggested the involvement of large scale brain networks beyond mere auditory and motor activity, such as visual areas, parietal cortices and areas of the default mode network, thus agreeing with previous group results [42].

However, the complexity and idiosyncrasy of musical improvisation might not be ideally captured using controlled experiments that compared improvisation with “non-improvisation” conditions, or mere effects of expertise in improvisation on brain plasticity, paradigms which are used in the vast majority of studies as shown in [8]. To address this bias, it has been argued that the study of the neuroscience of creativity, and in particular musical improvisation, would be better approached by setting up collaborations between scientists and artists in order to achieve both ecological and scientific validity [34]. Notably, a recent study performed EEG measurements on performers and audience in a live concert [18]. Their results demonstrate potential neural correlates of flow state using a measure of signal complexity, and this study more generally shows the feasibility of such ecological designs to better understand musical improvisation in a live context [18].

Building upon these different directions, our goal is to study musical improvisation from the point of view of an improviser, by implicating the musician in the scientific design of the experiment. We propose here to setup a collaborative process with the musician in order to define an appropriate paradigm, and repeated this paradigm in a series of rehearsals and public performances. Our objective is to maximize ecological validity by studying a single subject on many occasions, in an attempt at generalizing findings within this subject. By doing so, we also hope that such an approach can be of interest for the musician itself, by providing some scientific insights towards an introspection of his creative process.

The rest of this paper is organized as follows. In section 2, we describe our general setting, the collaboration with the artist, and the definition of an ecological paradigm to study musical improvisation. We performed EEG measurements on a musician during live concerts, followed by retrospective ratings of the performance. This paradigm has led us to consider a new hypothesis to test with regards to subjective time during musical improvisation. We present in section 3 a controlled paradigm design to test this hypothesis. Finally, we discuss our results and our approach in section 4.

## 2 Experiment 1 : ecological paradigm

### 2.1 Materials and methods

#### 2.1.1 Subject description

This study was performed on a single subject, also co-author of this manuscript, Christophe Rocher (CR), 53 years old. CR started playing the clarinets at the age of 7, and plays both the clarinet and the bass clarinet. CR has performed regularly in regional, national and international music scenes, in particular in the free improvisation scene, with ensembles of various sizes, as well as in performances with other artists such as dancers or spoken word artists. Importantly, the present study involves CR more than as a mere participation as a musician; we setup a collaboration with CR in order to define an appropriate approach to study musical improvisation from the point of view of an improviser. This collaboration was kept all along the project, but its goal was to assist on the definition of the main paradigm. As a consequence, the data collection performed in this study was agreed upon with CR in the preliminary phase of the study. An informed consent form was signed so that CR was aware what kind of data we were going to collect (EEG and audio), that he could decide to withdraw at any time, and that he could ask that his data was deleted at any time.

#### 2.1.2 Preliminary phase

We aimed at defining an ecological paradigm to study improvisation, with two aims. First, we tried to approach improvisation from the point of view of CR. The point made here consists in examining in detail the strategies developed by one particular improviser during his career, and document closely his creative process. Second, we target the study of subjective mental states associated with his performance. The proposed approach attempts at studying improvisation using a bottom-up approach, starting from the subjective experience of the improviser and in an ecological manner.

Experimental sessions consisted of free improvised concerts with an audience, followed by a relistening session. The goal of the relistening session was for CR to attempt a retrospective mental replay of his subjective experience during the performance. We aimed at documenting this retrospective phase. In preliminary experiments taking the form of private rehearsals, CR made an open commentary while listening to the performance. A first informal discussion around the content of these commentaries has enabled us to consider several emerging concepts : focus on improvisation, flow, satisfaction about the music being played, and the relationship between the musicians and subjective time perception. According to CR, these concepts were the ones that forge his everyday practice, and are related to the musical and personal objective occurring during a performance with an audience. At this stage in the project, we identified and acknowledged two important limitations in our approach. First, we were aware of the idiosyncrasies of these concepts, which may or may not apply to other professional improvisers. Second, as the open commentary of CR of his improvised performances tended to lean towards the same concepts, we decided to attempt a quantification of these concepts, by performing a continuous rating with three factors while listening to the performance.

Six rehearsals were performed in total, which are considered as the pilot phase of the project. During the first rehearsal, the retrospective phase consisted of the open commentary described above. During the second and third rehearsal, we asked CR to annotate the performance using a continuous rating with three factors. We agreed with CR on the meaning of the extreme values of these factors, and debriefed after each annotation session to make sure that the annotation were performed consistently.

The first factor was “focus”, and corresponds to how much CR felt he was successfully focused on improvising. A high value in Focus meant that CR was improvising while not being distracted. A low value meant that the focus on improvisation was compromised for various reasons. These reasons can relate to sonic or technical aspects of playing such as being in tune, having a nice clarinet sound, breathing. CR also reported higher level cognitive distractions related to the audience or music unrelated mind-wandering, in which case he also put a low value for focus. The second factor was “Subjective Temporal Resolution” (STR), corresponding to variability in the subjective quantization of time as retrospectively assessed by CR. According to CR, such a subjective quantization influences how he reacts to the music being played by other musicians, and can be loosely related to a subjective musical *tempo* (while the music itself often doesn’t have a clear tempo), or a clock with a period of a few hundred milliseconds up to a few seconds. Note that while STR is linked to the speed of subjective time, CR claims that it does not necessarily correspond to the speed of notes that he is currently playing, if he is playing at all, and we specifically tested this hypothesis (see section 2.3.4). CR reported that he consistently set low (respectively high) values of STR when his subjective quantization is slow (respectively fast). The third factor was “quality”, related to the personal satisfaction about the music being played. This factor judged *a posteriori* the quality of the music, from the point of view of CR, in terms of whether it corresponds to what he expects to offer to the audience.

These three factors were used for annotating the second and third rehearsal. The performances were annotated just after being played. A debriefing at the end of the third rehearsal was done and we agreed with CR that the third factor, “quality”, was most of the time highly correlated with “focus”, and it was also challenging to annotate three factors simultaneously and continuously while listening. We therefore decided to drop the “quality” factor. The three other rehearsals were used for piloting the EEG recording, getting familiar with playing with the EEG device while minimizing head or eye movements. We also performed these last rehearsals with a very limited audience (1 to 2 people) in order to have people actually listening, which according to CR helped him to be in a state closer to an actual performance.

### 2.2 Procedure

#### 2.2.1 Ecological paradigm

Here, we describe the final ecological setting that was used for the three public performances considered in this paper. Two performances took place in March 2019 in Brest, France, in front of audiences of 50 people (referred to as perf1 and perf2 in the rest of this paper). The third performance (perf3) took place in Montreal at the Montreal Neurological Institute in June 2019. Each performance was scheduled to last 20 minutes maximum, and the aim was to break it into two sessions of 10 minutes. The performances were followed by a 20 minute long talk and a discussion, presenting the project aims, and involving CR in the discussion with the audience. Following a preliminary recording session where we qualitatively assessed the impacts of blinks, eye movements and movement artifacts while recording and after recording, we discussed how to reduce them during the performance. It was agreed that CR would keep his eyes closed and that he would make his best to limit his movements (i.e body, fingers, breathing,…). A video of performance 3 can be found here https://www.youtube.com/watch?v=ILhaZYtW8fs, as well as a short documentary (in french) with excerpts of rehearsals and of performances 1 and 2, here https://www.kubweb.media/page/nautilis-neurosciences-jazz-improvisation-nicolas-farrugia/.

#### 2.2.2 Data acquisition

Each session was structured in the following way (Figure 1 panel a). CR played pieces in duet or trio lasting approximately 10 minutes. During each piece, we recorded audio and electroencephalography (EEG) on CR. Audio was recorded using a RME Fireface 400 FireWire audio interface, with two Neumann KM184 microphones. Microphones were placed to record the whole band for the subsequent relistening phase. The Bitwig software was used for the recording. EEG was acquired using an open BCI 8 channel Cyton amplifier. We used the headband kit to measure three frontal flat snap electrodes positioned on Fpz, Fp2 and Fp1, as well as two temporal dry comb electrodes located at FT7 and FT8. EEG was recorded at a sample rate of 250 Hz using the Fieldtrip buffer [40] and the EEG synth software (https://github.com/eegsynth/eegsynth). A one minute resting state was acquired, during which CR relaxed and prepared himself silently. This one minute resting state was part of the public performance and served as a silent introduction. CR was deliberately instructed to keep his eyes closed during the performances. Following each piece, CR listened back to the audio recording (no later than 24 hours following the performance), and performed the retrospective rating using the two factors Focus and STR, detailed in section 2.1.2. Retrospective rating was acquired using the Bitwig software using a USB-MIDI control interface with two continuous sliders.

**Figure 1:**
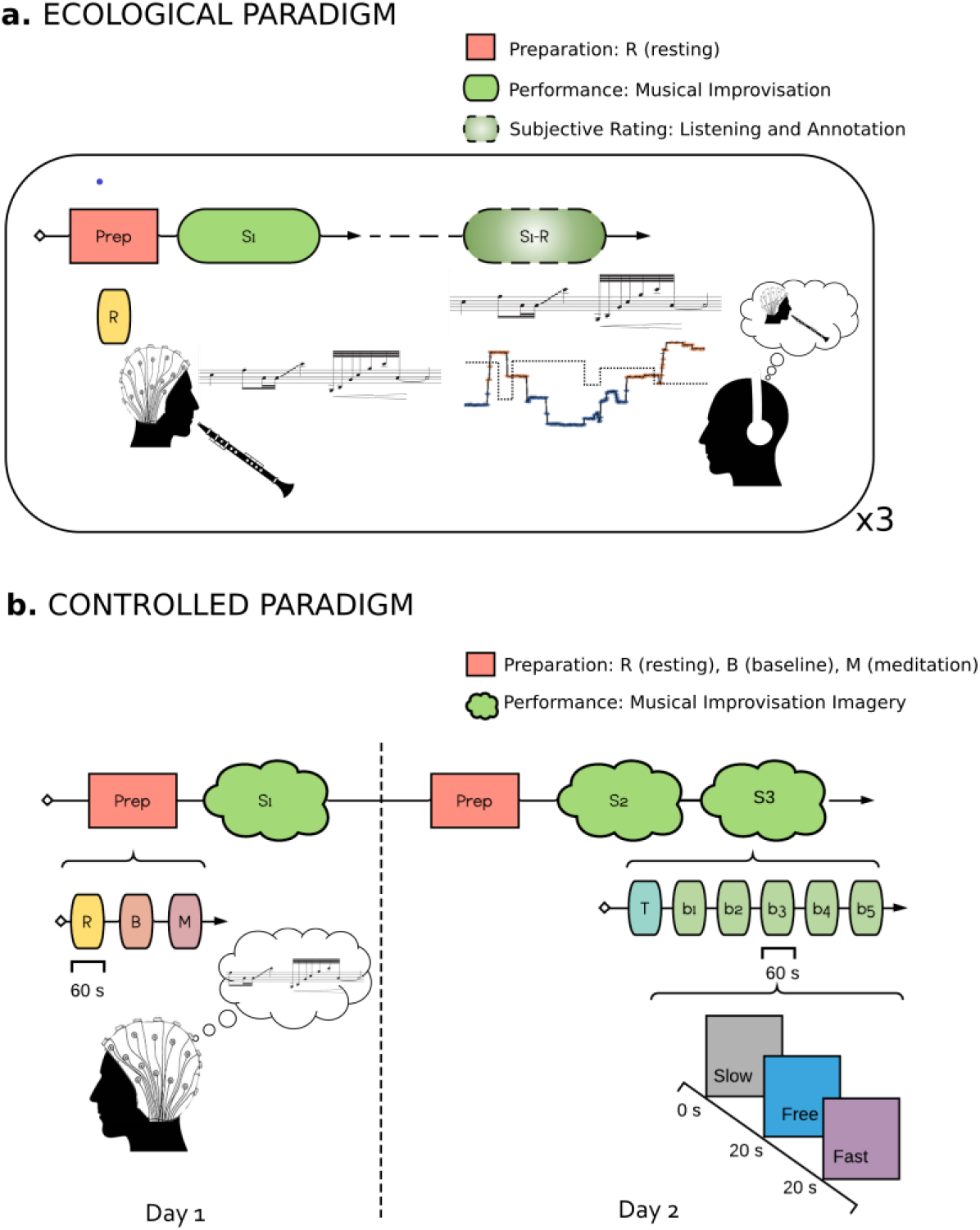
Experimental Protocols Schematics. a. Ecological Paradigm. The experiment included two parts. In the first, EEG was recorded while the subject performed musical improvisation. In the second part, the subject listened to its own performance and performed the retrospective rating using the two factors Focus and STR, detailed in section 2.1.2. b. Controlled Paradigm. The experiment was carried out in two days. In the first, the subject underwent a Preparation session where he performed 60 s of Resting (Eyes Opened), 60 s of Baseline and 60 s of Meditation. He then performed a musical improvisation imagery task with a Slow, Fast or Free conditions. The second part (two days later) was as the first, with the exception that two training sessions were performed.

### 2.3 Data analysis

#### 2.3.1 Behavioral data analysis

A qualitative analysis of the values taken by Focus, suggested that the Focus rating was generally high during performance (figure 2, right panel). Discussions with CR have led us to consider that Focus did not represent a source of variability inherent to musical improvisation, but rather was indicative of whether he reached the target state enabling him to improvise. As a consequence, in the rest of our analysis, we will only consider the STR rating. We used Hidden Markov Models (HMM) [46] to quantify the STR time series into discrete states. HMM is a probabilistic sequence model that estimates a series of hidden states from a set of observations. These hidden states are interpretable as causal factors of the probabilistic model (e.g. subjective “states” of STR). We considered a HMM with Gaussian emissions with two hidden states corresponding to *low* and *high* values of STR. We used the hmmlearn package (https://hmmlearn.readthedocs.io/en/latest/index.html) to learn the HMM model solving the iterative Baum-Welch Expectation-Maximization algorithm [15].

**Figure 2:**
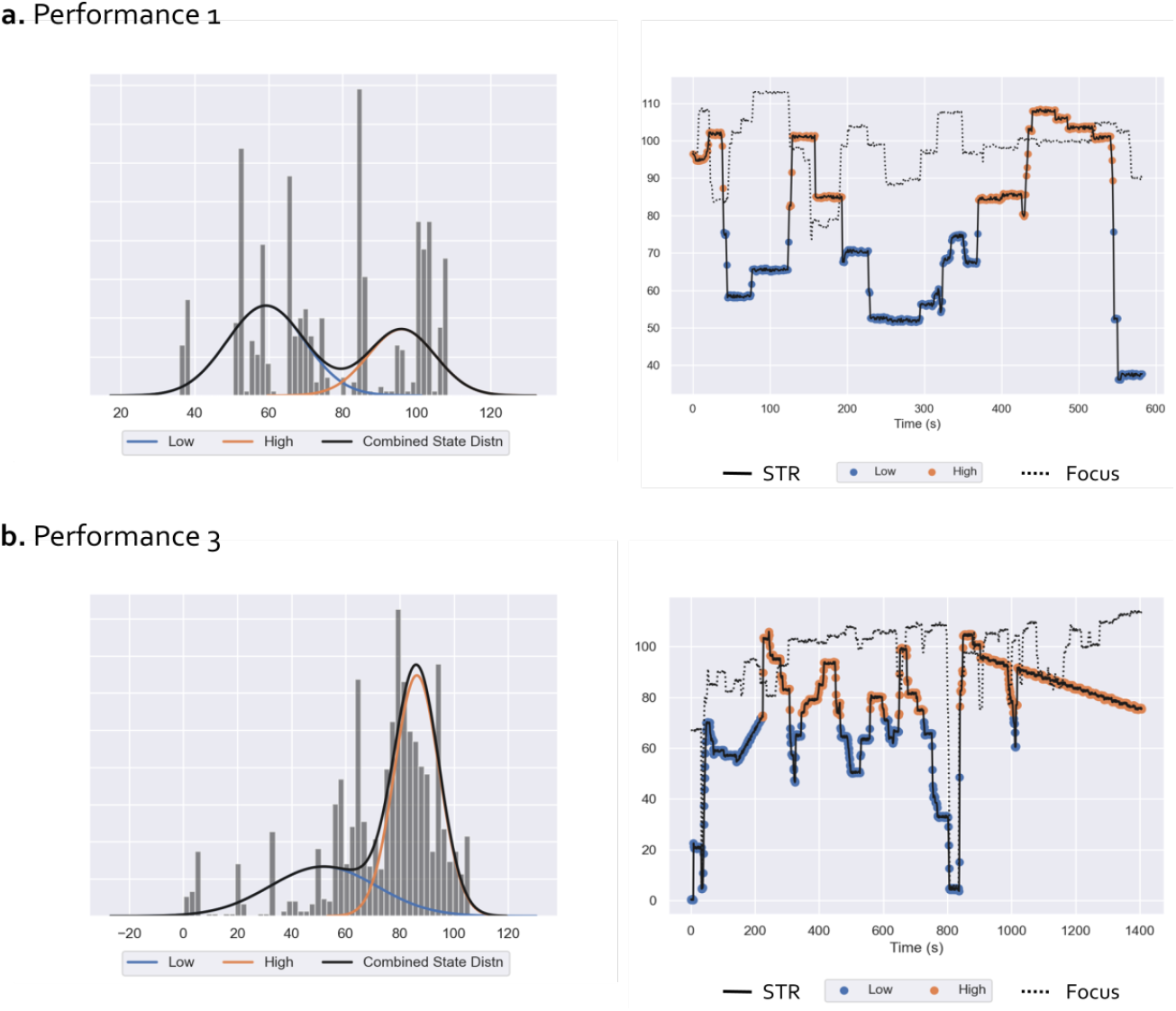
Ecological Paradigm: Results of HMM analysis of subjective rating scores for performance 1 (panel a.) and performance 3 (panel b.) Left: STR samples histograms and distribution (black) as a mixture of low (blue) and high (orange) states Gaussian distributions. Right: STR (solid line) and Focus (dotted line) time-series relative to the performances 1 and 3. For the STR, samples corresponding to the low and high states are labeled with blue and orange markers respectively.

#### 2.3.2 EEG preprocessing

EEG data were preprocessed using the MNE-python toolbox [23]. First, Signals were bandpassed filtered with a FIR (Finite Impulse Response) filter in the 1 - 40 Hz frequency band. To reduce eye movement artefacts, we perform Independent Component Analysis (ICA) using the *fastica* algorithm [25] applied to continuous data. We ran an autodetection algorithm to find the independent component that best matched the ‘EOG’ channel (prefrontal electrode Fp2). ICA components that strongly correlate with the EOG signal were then removed (adaptive Z-score threshold = 1.6) and the EEG signal was reconstructed with the remaining components. In order to reject residual movement artifacts, we then segmented data into consecutive epochs of 3 seconds and remove those in which the signal amplitude of one or more channels exceeded a threshold set to keep the 85% of recordings.

#### 2.3.3 Time-frequency analysis and decoding model

We performed a time-frequency analysis using multitaper filters to estimate the EEG power spectral density and the average power in different frequency bands (theta, alpha and beta) computed with reference to the individual alpha frequency [4] of the subject (IAF= 9.3 Hz). Based on IAF frequency we estimated the theta, alpha and beta bands respectively equal to [4.5-7.5] Hz, [7.5-11.5] Hz, [11.5-25] Hz. We estimated the EEG power for 3 seconds epochs and assessed whether it could predict the STR as being low or high using a decoding model with a Support Vector Classifier (SVC) and a radial basis function kernel with regularization (*C* = 1, penalty on the squared *l*_2_ norm), implemented in the scikit-learn package [41]. In order to test for within-sample generalization of our decoding model using the data at hand, we used a stratified K-fold cross-validation with 4 folds in order to consider the same percentage of samples of each class per fold. We measured classification accuracy and f1-score for each class and fold. In order to provide an even more conservative robustness assessment of our results, we performed a hundred repetitions of the same cross-validated SVC training using random permutations of class labels (see https://scikit-learn.org/stable/modules/generated/sklearn.model_selection.permutation_test_score.html). This permutation test score provides an estimation of the chance level of our decoding model according to the variance in the dataset. We performed a post-hoc univariate statistical inference analysis by investigating changes in the different frequency bands related to the STR state. More specifically, we assessed differences between average EEG power during low and high states in the theta, alpha and beta bands by means of a pair-wise two-sided Welch t-test using the SciPy Stats library (https://docs.scipy.org/doc/scipy/reference/stats.html#module-scipy.stats).

#### 2.3.4 Audio analysis

The aim of the audio analysis was to characterize note density (number of notes played every second) and volume of the musical performance of CR on the three performances, in order to study their relationship with Subjective Time Resolution (STR) and EEG rhythms. This analysis was performed separately for each performance. First, we separated the different musical instruments recording before counting notes. For each performance there were three instruments: clarinet (CR), drums and double bass for the first two performances and clarinet, double bass and trumpet for the third performances. As we are focusing on the performance of CR (specifically, the number of notes he played) we separated the clarinet from the two other instruments. We used the 4 stem source separation pretrained model from the library Spleeter [24, 53] developed and open-sourced by Deezer. Spleeter allowed us to split music instruments leveraging pre-trained neural networks implemented in TensorFlow. Next, note onset detection was performed on the clarinet track separated during step 1. For this, we used an onset detection method available in the librosa package (https://librosa.org/doc/main/generated/librosa.onset.onset_detect.html#librosa.onset.onset_detect) that works by detecting local peaks in the envelope strength, using hyperparameters set on a large music database. We validated note density extraction qualitatively by retrieving several samples for each note density bin and checked qualitatively if the note density was well estimated. Then, we assessed the link between note density during the musical performance and subjective state (STR). First, we estimated the distribution of note density values across performances. As there are many moments during which CR does not play, we get an important proportion of note density values of 0. However, non-zero density values followed an approximately gaussian distribution, that we binned in three categories including values b. We tested the hypothesis whether CR plays more notes per second as a function of subjective state (high or low STR), by using the annotations of the HMM (see behavioral data analysis). We also compare qualitatively the count of note densities equal to zero (i.e. moments during which CR does not play for one second) in the subjective states. We also tested the hypothesis that the volume of musical performance was related to subjective state. To this end we calculated the RMS energy of each performance using the librosa function librosa.rms() and compared RMS distributions for low and high STR using Welch’s t-test.

Finally, to test whether note density had a significant impact on EEG power in different frequency bands we used the four above mentioned categories: silence (note density equal to zero), low density, medium density and high density, and computed alpha, beta and theta EEG power in each category. We tested for associations between power and note density using a Kruskall Wallis test, and performed pairwise t-tests for post-hoc comparisons.

### 2.4 Results

#### 2.4.1 Analysis of subjective ratings

Results of the HMM analysis of STR time-series for performances 1 and 3 are reported in Figure 2. Two hidden states corresponding to low and high STR values were identified: the relative estimated Gaussian distributions are represented in the left panels of Figure 2 while their values during the performance, together with the Focus index trends are reported in the right panels. EEG recordings of perf1 were highly contaminated by environmental and movement artifacts (see EEG results section). Since we only examined behavioral indexes relative to preprocessed EEG epochs, the number of samples of STR and Focus for perf1 is drastically reduced as compared to perf3, resulting in a sparser histogram distribution and shorter time-series.

We note that Focus values are generally staying high during performance, with a few disrupted moments occurring with low values. As a consequence, we did not model the variability in Focus, and the rest of the analysis was performed with respect to HMM states obtained by the analysis of STR values.

#### 2.4.2 Link between subjective states and audio-derived features

We first sought to test whether the identified subjective states (low and high STR) were related to the volume of the overall performance. RMS energy (volume) was lower in the low STR state than in the high STR state for perf2 (t(1046)=−7.7, p = 1e-14), while it was higher in low STR than in high STR state for perf1 (t(581)=7.0, p=5e-12) and perf3 (t(1465)=10.9, p=1e-26). Next, we analyzed the note count played by CR per second (note density) as a function of subjective state. We found quantitative differences when examining moments during which CR does not play (approximated by a note density of zero, meaning no note played for one second), with a lower number of zero densities in low STR compared with high STR in performance 3 (3 for low STR, 29 for high STR), while the opposite was true for performance 1 (47 for low STR, 16 for high STR) and performance 2 (104 for low STR, 54 for high STR). Finally, slightly more notes per second were played in low STR than in than STR for performance 2 (mean density for low STR = 11.1 note per second, 9.8 for high STR, t(888) = 5.7, p = 1e-8) and performance 3 (mean density for low STR = 10.3 note per second, 9.4 for high STR, t(1367)=4.8, p = 1e-6), but no difference was found in performance 1 (mean density for low STR = 8.7 note per second, 8.6 for high STR, t(518) = −0.4, p=0.7). A graphical depiction of those results for performances 1 and 3 is given in supplementary Figures 1 and 2, panel a.

#### 2.4.3 Link between EEG oscillatory power and STR

A hardware problem with the EEG amplifier occured when recording perf2, so we only report results on perf1 and perf3. The EEG recordings of perf1 being very noisy, only the equivalent of 10 minute recordings survived artifact rejection and were considered for further analysis. For perf1, SVC results indicated that high and low STR states could be classified with an average accuracy of 0.69 ± 0.11 (standard deviation across folds) (f1-score high 0.63 ± 0.16 - 94 examples-, f1-score low 0.74 ± 0.10 - 87 examples-). Similarly for perf3, a SVC trained on EEG power distinguished low from high states with an average accuracy of 0.69 ± 0.11 (f1-score high 0.69 ± 0.16 −165 examples-, f1-score low 0.66 ± 0.12 −170 examples-). The permutation test in both cases indicated that the decoding model performed significantly better than chance (*p* < 0.01)

Post-hoc statistical analysis (Figure 3) for the different frequency bands revealed that theta, and beta average power was higher in the high STR state condition as compared to the low condition both in perf1 (theta: p=2.5e-08, t(155)=−6.30; beta: p=6.4e-07, t(139)=−5.69) and perf3 (theta: p=3.1e-06, t(239)=−5.24; beta: p=1.9e-05, t(226)=−4.8). This trend was also observed in the Alpha band for perf1 (p=1.8e-07, t(154)=−5.91) but was not significant in perf3 (p=7.8e-01, t(201)=−1.7).

**Figure 3:**
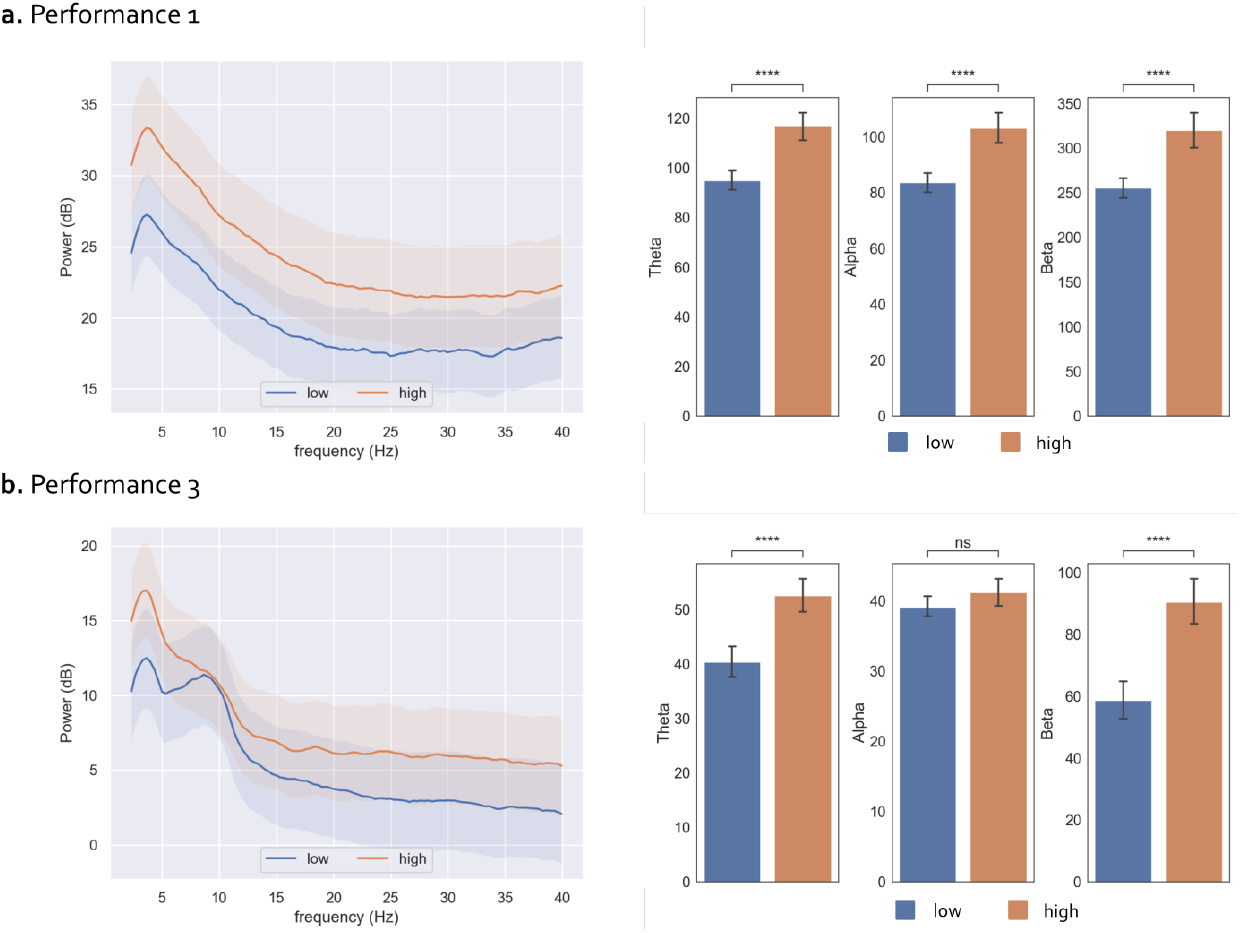
Ecological Paradigm: Results of EEG post-hoc frequency analysis in relation to STR for performance 1 (panel a.) and performance 3 (panel b.) Results represent EEG power averaged across electrodes. Left: EEG Power Spectral Density (mean across 3 s epochs) corresponding to low (blue) and high (orange) states, with 95% confidence intervals. Right: Bar plots representing the EEG power (mean ± std) in the Theta, Alpha and Beta bands in low and high states. Square brackets indicate significant differences as assessed with pair-wise two-sided Welch t-test (**** *p* < 0.00001)

#### 2.4.4 Link between EEG oscillatory power and audio-derived features

The volume of performance 1 was positively correlated with theta power (*ρ* = 0.2, p=0.03), but not with alpha (*ρ* = 0.09, p=0.3) and marginally with beta power (*ρ* = 0.2, p=0.07). Interestingly, the opposite relationship was found for performance 3, with volume being negatively correlated with alpha power (*ρ* = −0.2, p=0.007), beta power (*ρ* = −0.2, p=4e-4) and theta power (*ρ* = −0.3, p=1e-5). We subsequently tested for associations between note density and EEG power, by using four bins of note density (zero, low, medium and high) in separate Kruskal Wallis tests for each frequency band. In performance 1, we found a significant effect of note density on alpha power (*t* = 9.5, *p* = 0.02), beta power (*t* = 9.2, *p* = 0.02) and theta power (*t* = 11.3, *p* = 0.01). A similar pattern of results was obtained for performance 3, with an effect of note density on alpha power (*t* = 11.6, *p* = 0.001), beta power (*t* = 8.8, *p* = 0.03) and theta power (*t* = 10.8, *p* = 0.01). In performance 3, post-hoc t-test revealed higher EEG power during zero note densities than during the three other levels of note density, with the largest effect obtained in the three frequency bands (all comparisons between zero density and other levels with *p* < 0.01). In performance 1, lower alpha and beta power was observed during zero note density than during medium note density (*p* < 0.05). Lower theta power was also observed during zero density than during low (*p* = 0.004), medium (*p* = 0.003) and high (*p* = 0.04) densities. These results are illustrated in supplementary figures 1 and 2, panel b.

## 3 Experiment 2 : controlled paradigm

### 3.1 Materials and Methods

#### 3.1.1 Procedure

The experimental paradigm is described in Figure 1 panel b. The main goal of this experiment is to manipulate STR in a controlled setting, by asking CR to perform a musical improvisation imagery task, while constraining himself to stay in a particular state with respect to STR, thus corresponding to different quantization of subjective time. Three conditions were considered : Slow, Fast and Free. Slow and Fast corresponded respectively to slow or fast quantization of subjective time. These conditions are considered according to the states found when analyzing the retrospective rating phase of Experiment 1 (see sections 2.1.2 and 2.4.1). For these two conditions, the instructions given to CR were to imagine he was improvising while keeping a subjective state that he would have rated as either Low or High STR during the retrospective phase. The third condition, Free, corresponded to musical improvisation imagery without constraints on a subjective state related to STR. The experiment was carried out over two separate sessions on two different days. During each session, we performed a preparation phase which consisted of a one-minute long resting state with eyes open (R), a one-minute active baseline consisting in counting backwards (B), and a one-minute meditation phase (M) during which CR attempted to focus on breathing. In both B and M phases CR kept his eyes closed. These conditions were implemented in order to have clear cut comparisons between states with different mental workload in order to check signal quality, and were not analyzed further (except for the B condition which was used to determine IAF). Following the preparation phase was the musical improvisation imagery task. The experiment was organized into a training block, followed by 5 blocks. The order of conditions was randomized and counterbalanced across blocks, and each condition was presented fifteen times in total. Each block consisted of three consecutive trials of twenty seconds. Instructions were given vocally at the beginning of each trial, with the experimenter pronouncing the words “Slow”, “Fast”, or “Free”. These instructions were explained before the training block. A debriefing after the practice block of each session was made, in order to gather informal feedback on the feasibility of the task. Within a block, a condition might be repeated, in order to avoid that CR predicts the third condition and change his strategy accordingly. A short break was done after each block. During the first session, we performed only five blocks, while two times five blocks were done during the second session, with a longer break between after the fifth block.

#### 3.1.2 EEG acquisition

The measurements were done in two slightly different settings for day 1 and day 2. During day 1, we performed the experiment in a moderately quiet environment, a common space with a few people passing. During day 2, we performed experiments in a quiet room with only the experimenter and CR. As for the ecological paradigm, at the beginning of each session CR was given precise instructions to keep his eyes closed during musical imagery. CR performed all the conditions while closing his eyes, and could open his eyes between blocks. CR was sitting in front of a white wall with the experimenter in his back. EEG was acquired using the same amplifier and software setup than in Experiment 1 (see section 2.2.2), but with a different electrode montage. Four goldcup electrodes were positioned at O4, P4, C4 and Fp4 using conductive paste.

### 3.2 Data analysis

#### 3.2.1 EEG preprocessing

As for the first experiment, we performed ICA on the band-pass filtered EEG signals (1.0-40.0Hz) in order to reduce eye movements artifacts using the prefrontal electrode Fp2 as a proxy for the EOG channel. We then divided each block into 20 seconds segments according to the trial onsets, and removed the first 5 seconds of each trial to reduce the effect of transition between trials. Finally, trials were segmented into consecutive epochs of 3 seconds, and epochs in which the signal amplitude of one or more channels was high were removed, using a threshold set to keep 90% of data.

#### 3.2.2 Statistical analysis

Individual Alpha Frequency (IAF) [4] was determined by finding the individual dominant EEG frequency in the baseline signal. As for the first experiment the resulting frequency bands were: theta [4.5-7.5] Hz, alpha [7.5-11.5] and beta [11.5-25] Hz. To conduct our analysis, we estimated the average power spectral density across the four electrodes (Fp2,C4, P4, O2) using multitaper filters, and we computed the power in the different frequency bands. The 3 second-long epochs were labelled with the corresponding condition (free, slow, and fast) and Welch pair-wise t-tests were performed to assess the effect of condition on the EEG power magnitude in different frequency bands of interest. Results were corrected for multiple comparison according to the Bonferroni correction.

### 3.3 Results

#### 3.3.1 Behavioral results

CR indicated that he could generally perform the task, and gave details about specific mental imagery strategies that he used to help him perform the task correctly. CR indicated that he imagined himself playing in specific places, with specific people. As a consequence, the feedback given by CR suggest that he engaged more than in a constrained mental imagery exercise.

#### 3.3.2 EEG results

Statistical analysis results (Figure 4) revealed that beta power was higher in the Slow condition as compared to the Fast condition (p=0.049, t(116)= 2.83). The Free condition was associated with a higher beta (p=0.011,t(119)=3.31)) and theta power (p=0.008, t(134)=−3.41) if compared with the Fast condition. This trend was also observed in the Alpha band but did not survive Bonferroni correction.

**Figure 4:**
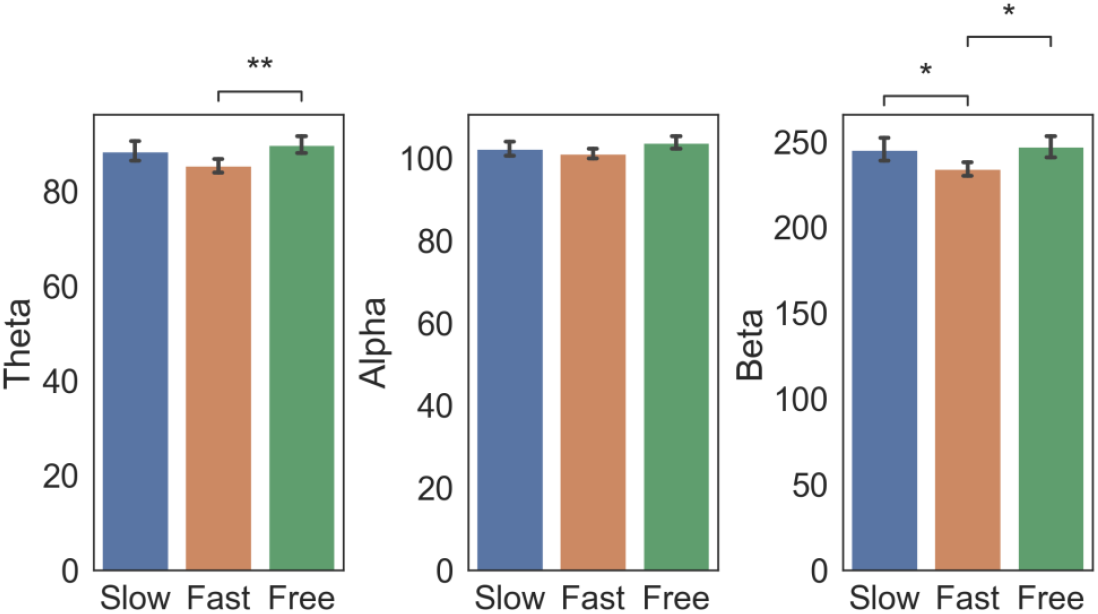
Controlled paradigm: EEG power changes as a function of subjective time condition. EEG power (mean ± std across electrodes Fp2, C4, P4 and O2) in the Theta, Alpha and Beta bands in epochs corresponding to Slow, Fast and Free condition. Square brackets indicate significant differences as assessed with pair-wise Welch t-test (* *p* < 0.05, ** *p* < 0.01, Bonferroni corrected)

## 4 Discussion

### 4.1 Summary

We have presented an ecological paradigm of musical improvisation live performance with an audience, consisting in EEG measurements of an improviser, followed by a listening phase with retrospective rating. The objective of the rating was to perform *a posteriori* mental replay of the subjective state of the performer. A discussion with the improviser led us to consider two continuous factors when rating performance: Focus and Subjective Temporal Resolution (STR). The meaning of these factors was discussed, piloted and consistently confirmed with the subject. Focus measured a general tendency to “feel in the music”, or “being in the zone”. STR measured subjective temporal resolution, which indicates whether the improviser was in a state of slow or fast subjective time quantization. Using a decoding model trained on EEG power during performance, we found that states of high and low STR could be reliably distinguished, and were related to increases in average theta and beta power during the high STR state. We also showed that CR played more notes in states of low STR than in states in high STR, in performances 2 and 3, and that states of low STR could be associated with higher volume (resp. lower volume) than states of high STR in performances 1 and 3 (resp. performance 2). When testing for associations between EEG power and number of notes played, we found global changes in power when CR plays compared to when he does not, but the number of notes played did not influence EEG power. We also found that EEG power was weakly correlated with volume in performance 1, and negatively correlated with volume in performance 3. In a second experiment in a controlled setting, we designed a musical improvisation imagery experiment targeted at testing differences in brain oscillations with respect to STR, and we found elevated EEG power in the beta band when CR was in a subjective state of low STR.

### 4.2 Musical improvisation as a target subjective state ?

We approached the question of characterizing improvisation as a target subjective state, measured by two factors in a retrospective rating. The concept of musical improvisation as a subjective state was previously proposed [18, 31], and was interpreted in the context of flow state [13]. In the following, we attempt to interpret the two factors we measured, Focus and STR.

What we have termed Focus in this study corresponds to a component of a common definition of flow state, “the holistic sensation that people feel when they act with total involvement,” [13], and has been extensively studied, including in the music improvisation literature (e.g. see [10, 18, 32, 54]. Previous research on flow state during improvisation was mostly done using interviews and observations [54]. In our case, a qualitative analysis of the values taken by the Focus rating, together with informal observations discussed with the performer, suggested that Focus was generally staying high during musical improvisation performance, and corresponded to a target for appropriate performance. Preliminary exploratory correlation analysis between EEG power and the Focus factor did not reveal any link in our measurements.

On the contrary, STR, considered by CR as the quantization of subjective time, has not been previously documented as an aspect of flow state. Previous studies have proposed that the distortion of subjective time perception is an important part of the psychological state of flow [13,14]. Such an account usually refers to the feeling of an accelerated passing of time during flow state, and has been measured previously in laboratory conditions [26], as well as in previous studies such as in gaming [39] and music performance [10]. To our knowledge, STR has not been a measure of interest in previous studies on flow state of musical improvisation. We therefore have to turn to the temporal processing and the attention literature to bring some light on our findings.

### 4.3 Subjective temporal resolution and temporal processing

Temporal resolution has been measured experimentally with simultaneity judgment tasks [52], in particular audiovisual simultaneity. Recent reviews have shown considerable variation in task performance according to stimulus modality, inter-individual differences, age, as well as subjective states [2], [63]. Interestingly, musical training has been shown to influence audiovisual simultaneity judgments [27], suggesting that long-term training modulate musician’s ability to integrate audiovisual information concurrently. Recently, audiovisual simultaneity has been linked to phase resetting in the EEG beta band [28]. However, we cannot comment on whether such integration processes are related to our findings on STR, as simultaneity judgments can only be done in lab settings with controlled stimuli. Another account related to temporal resolution is the concept of temporal receptive windows [30], with a hierarchy spanning from early auditory cortices at the smallest time scales (less than one second), up to parietal and frontal areas for the largest ones (up to a minute). Here, our attempt at measuring STR had the objective of tapping into subjective processes related to the quantization of subjective time. While a very large body of literature exists on subjective timing paced by an internal clock with periods from seconds to minutes (as initially proposed by [11], see [1] for a recent review), we are interested here in shorter periods in the range a few hundred milliseconds up to a few seconds. Such short time scales have been tackled in studies on the neural basis of temporal processing.

### 4.4 Brain oscillations and subjective temporal resolution

The proposed STR measure as well as our EEG results may also be interpreted with regards to a large body of work on electrophysiological correlates of temporal processing [33, 57], in light of predictive processes (such as isochronous sounds or beat perception), duration estimation and attention to temporal events [38]. We note first that no single EEG frequency band has been dominantly associated with temporal processing, as comprehensively shown in the cross-study review by [57]. More specific effects have been suggested in different types of paradigm. First, it has been shown that temporal expectations may modulate power in the theta band, as well as the coupling between theta phase and beta power [12], which could indicate the existence of a central mechanism for controlling neural excitability according to temporal expectations. These results have been recently complemented by a study that combined electrical stimulation and reanalysis of previous EEG data, showing an intrinsic role of beta oscillations in the memory of temporal duration [58]. The beta band has also been associated with effects of temporal prediction in the case of beat-based timing in perception [22] and imagery [21]. Finally, a classical paradigm to study temporal attention consists in providing a cue that predicts (or not) a short or long foreperiod between a warning stimulus and an imperative stimulus requiring a motor response. This paradigm revealed shorter reaction times when the cue successfully predicts the length of the foreperiod, together with an increases amplitude of the Contingent Negative Variation [35], as well as an increased EEG power between 6 and 8 Hz for stimuli with short foreperiods compared to long ones [3]. These results suggests that the brain allocates a temporal attention window of variable length mediated by underlying oscillatory mechanisms, namely the magnitude of EEG power in the 6 to 8 Hz band (upper theta band).

In experiment 1, we found a higher power in low frequency oscillations (4.5 to 7.5 Hz, dubbed theta in our study) and beta band (11.5 to 25 Hz) with high STR compared to low STR. As we are associating a retrospective subjective rating with EEG acquired in the presence of noise and movement, we attempted at disentangling the effect of overall volume and quantity of motor commands (approximated by a measure of note count per second) on the measured EEG. In performance 1, we found a weak correlation between theta power and volume, as well as a global increase in EEG power when CR plays compared to when he doesn’t. The opposite pattern was found in performance 3, with a negative correlation between volume and EEG power, as well as a global reduction of EEG power when CR plays. However, in both performances, the number of notes played did not influence EEG power. Importantly, we also showed using audio analysis of performances 1 and 3 that high STR was associated with lower note counts and higher volume than low STR. Therefore, it is unlikely that the EEG modulations we observed with respect to STR are solely due to motor activity, as otherwise we may expect to observe a relationship between EEG power and note count. Finally, we remark that the effect of STR on EEG power was consistent in both performances. This suggests that STR as measured in this ecological paradigm might reflect an underlying endogenous timing mechanism that calibrates the duration of a temporal window of integration, or equivalently, the rate of a sampling mechanism involved in musical performance and perception. This interpretation would fit with the description of the behavioral relevance of STR as discussed with CR during the definition of the protocol. It is obviously difficult to compare the ecological paradigm of experiment 1 with controlled experiments such as the ones mentioned previously, as we do not have controlled stimuli and multiple repetitions.The choice of performing a first experiment in an ecological setting was essential to define behavioral indexes related to the subjective experience of the musician, but came with some drawbacks. The main one is the limited quality of EEG signals collected in an environment exposed to noise and while CR was performing (e.g. freely moving). This compromised EEG recording during perf 2 and affected perf 1 signal quality. These limitations also motivated us to perform a second study in a controlled setting, where we could experimentally manipulate the subjective time state and assess STR changes on good-quality EEG recordings. As a consequence, we attempted to test specifically the effect of varying the rate of this sampling mechanism by defined a controlled paradigm. In experiment 2, we instructed the subject to perform musical improvisation imagery while keeping a specific state of STR. In a third condition, no constraint was given and the subject could perform imagery without keeping a constant STR state. We found an elevated theta and beta power when comparing the Free (unconstrained) condition with the Fast condition (corresponding to a high STR state), as well as higher beta power for Slow compared to Fast. While it can seem surprising to find a reverse effect than in experiment 1, it is difficult to conclude as theta and beta power was overall higher in the Free condition, which is the one that is closer to the ecological paradigm. Nevertheless, our results suggest that oscillatory power in the theta and beta band is correlated with an internal, subjective temporal processing system related to STR.

### 4.5 Brain oscillations and flow state in musical improvisation

A qualitative analysis of our ecological paradigm led us to consider the first rating, Focus, as an indicator of flow state during improvisation. We did not find any statistical association between the values of Focus and EEG power spectrum. However, in experiment 2, we did find a higher power in the theta and beta band when comparing the Free condition with the Fast condition. In this experiment, the Free condition corresponded to an unconstrained, more natural situation with respect to experiment 1, in contrast with the Slow and Fast conditions that instructed CR to perform mental imagery of a specific STR state. Therefore, the power increase observed in the Free condition may be interpreted in light of previous findings that showed EEG activity increases when comparing improvisation with “non-improvisation” [9,49]. Note however that the observed power increase might also be interpreted in a more general framework of creativity and flow state. Several studies have suggested a correlation between alpha-band activity and creative tasks [51]. Generally, it has been observed that tasks requiring greater creativity resulted in higher alpha power [20]. In particular, musical improvisation studies have reported higher alpha power in central and posterior regions of the brain, and a deactivation in prefrontal regions during the experience of flow [16]. Overall, the majority of the studies investigating creativity and musical improvisation report changes in alpha power, some studies even report clearer changes specifically in upper alpha [9, 49]. In experiment 2, the power increase between Free and Fast was found in the upper frequency band [11.5-25] Hz, as we defined alpha as [7.5-11.5] Hz in which only a trend towards statistical significance could be observed. As a consequence we can situate our results among previous studies, while keeping it clear that we only considered one expert subject. This effect requires replication with a larger and more diverse sample, and could be the goal of future controlled studies attempting at examining musical improvisation or creativity using mental imagery.

### 4.6 Implications for the artistic endeavour

The proposed collaboration between arts and sciences represents an original contribution towards artists in terms of imagination, a resource for them to explore new ideas. Personal introspection in the form of retrospective ratings has the potential to give artists a special insight into creation and musical practice. Open questions arise with respect to understanding the link between subjective states and musical outcomes, and such an understanding could potentially enhance the creative process. Furthermore, the discovery of experimental research and neuroscientific methods could bring artists with several new insights. Such collaborations could help make the artists aware that the scientific view of artistic creation contribute to a better understanding of creativity [34]. Such an endeavor may challenge the place of the musician as part of a complex, dynamical system including the other musicians and the audience. This questioning is in line with recent accounts on understanding musical creativity using the embodiement framework and dynamical systems [55]. Another contribution for artists is to learn about new technologies available today, with the idea of possibly directing musical and technological research towards the fabrication of new tools for musical computing, using for example neurofeedback or the sonification of brain waves. The wealth of research on brain computer interfaces, neurofeedback [50], and music information retrieval [36], could potentially contribute to the future of musical creation.

### 4.7 Limitations and perspectives

The limitations in this study are mostly inherent to the choices made regarding the ecological setting and the collaboration with a musician. As we considered a single subject, we do not have clear indications on the ability to generalize the concepts developed here and the findings to other musicians or other creative process. Future studies could attempt at testing hypothesis related to STR or flow in ecological settings using larger groups of musicians. In addition, while we decided early on to focus on a single subject, we relied only on retrospective reports and EEG recordings. The use of retrospective reports is limited by the metacognitive abilities of the rater, namely his ability to perform mental replay of the improvised performance. Such an ability might not be present with all musicians, which is another limitation towards a generalization of this procedure. Alternatively, future studies could consider semi-structured interviews in addition to retrospective ratings, which could potentially alleviate the bias introduced by ratings, while giving a richer qualitative view on the creative process, as done in previous musical improvisation studies [54]. Another concern in our study is the lack of control for the content of musical imagery in experiment 2, in particular regarding the number of notes imagined. This could have an effect on resulting oscillatory power, and we plan to design future controlled experiments to try to disentangle imagined content and subjective state. Finally, as we measured brain activity on a single subject using EEG during musical performance, the measured signal is largely contaminated with movement artifacts and other sources of noise inherent to the ecological context. While we attempted at controlling for motor commands associated with clarinet playing by estimating note density, we cannot easily rule out the presence of other cognitive mechanisms, such as the attention to other players, auditory working memory, or simply cognitive load, that might influence the measured EEG in such a free, naturalistic setting. The setup used in this study is simple and lightweight, but it includes too few electrodes to enable a high resolution study of the complex mechanisms involved in naturalistic musical improvisation. Future studies could build upon this study, and attempt to measure brain activity in ecological settings with a higher resolution. In addition, one way to limit contamination by movement artifacts would be to consider using functional near infrared spectroscopy (fNIRS) and motion capture simultaneously with EEG in order to provide a complementary view on brain activity while accounting for movement.

### 4.8 Conclusion

In this study, we have setup a collaboration with an artist, CR, performing free musical improvisation. This collaboration has led us to define an ecological paradigm to study musical improvisation during live performances with audiences, using retrospective ratings and electroencephalography. We have suggested a measure of Subjective Temporal Resolution as a correlate of a subjective state related to the quantization of internal time of the improviser during performance, and were able to relate this measure to EEG oscillatory power in the theta / low alpha and beta band. We subsequently devised a controlled musical improvisation imagery experiment and found a relationship between constraints on subjective time and oscillatory power in the EEG. Our results bring an original perspective on the study of musical improvisation and creativity, by showing the potential of single subject studies and ecological paradigms.

## Author Contributions

NF and CR designed the study and ran the experiments. AL, GL, JB and NF analyzed the data. All co-authors contributed to writing the manuscript.

## Acknowledgments

We’d like to thank Andrea Schiavio for his support and encouragements in the early phase of this project. In addition, we would like to thank the city of Brest, Brain Awareness Week and the Society for Neurosciences for enabling us to play the first two performances, as well as the Montreal Neurological Institute, and the Convergence Initiative, who supported our performance in Montreal. Finally, we d like to thank the reviewers, whose critical feedback has greatly improved the manuscript.

## Data Availability Statement

The code and data for this study can be found at the following url https://github.com/alixlam/Brainsongs1.

