## Supplementary figures and images for "Beta and theta oscillations correlate with subjective time during musical improvisation in ecological and controlled settings: a single subject study"

### Supplementary Figure 1

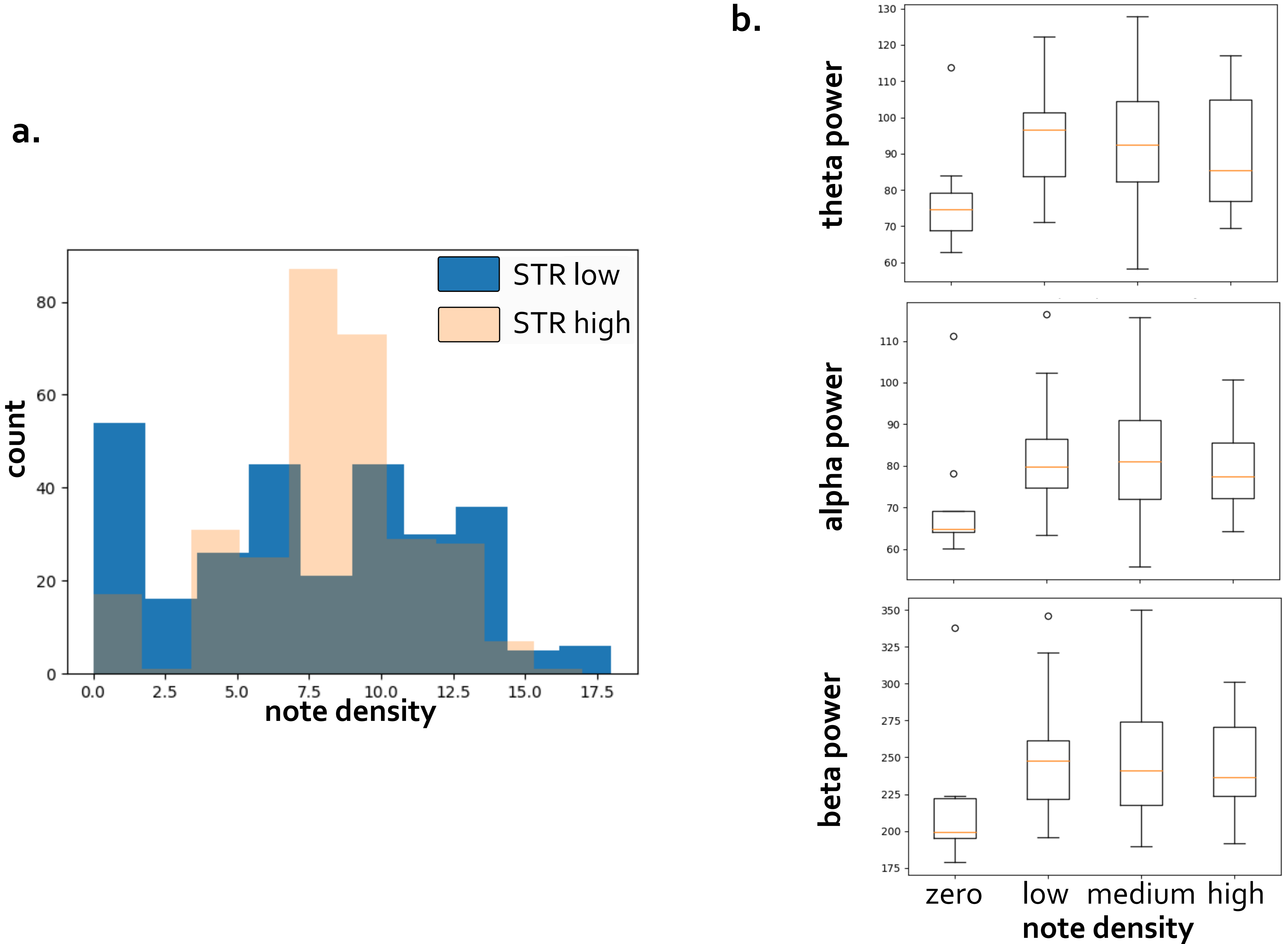

### Supplementary Figure 2

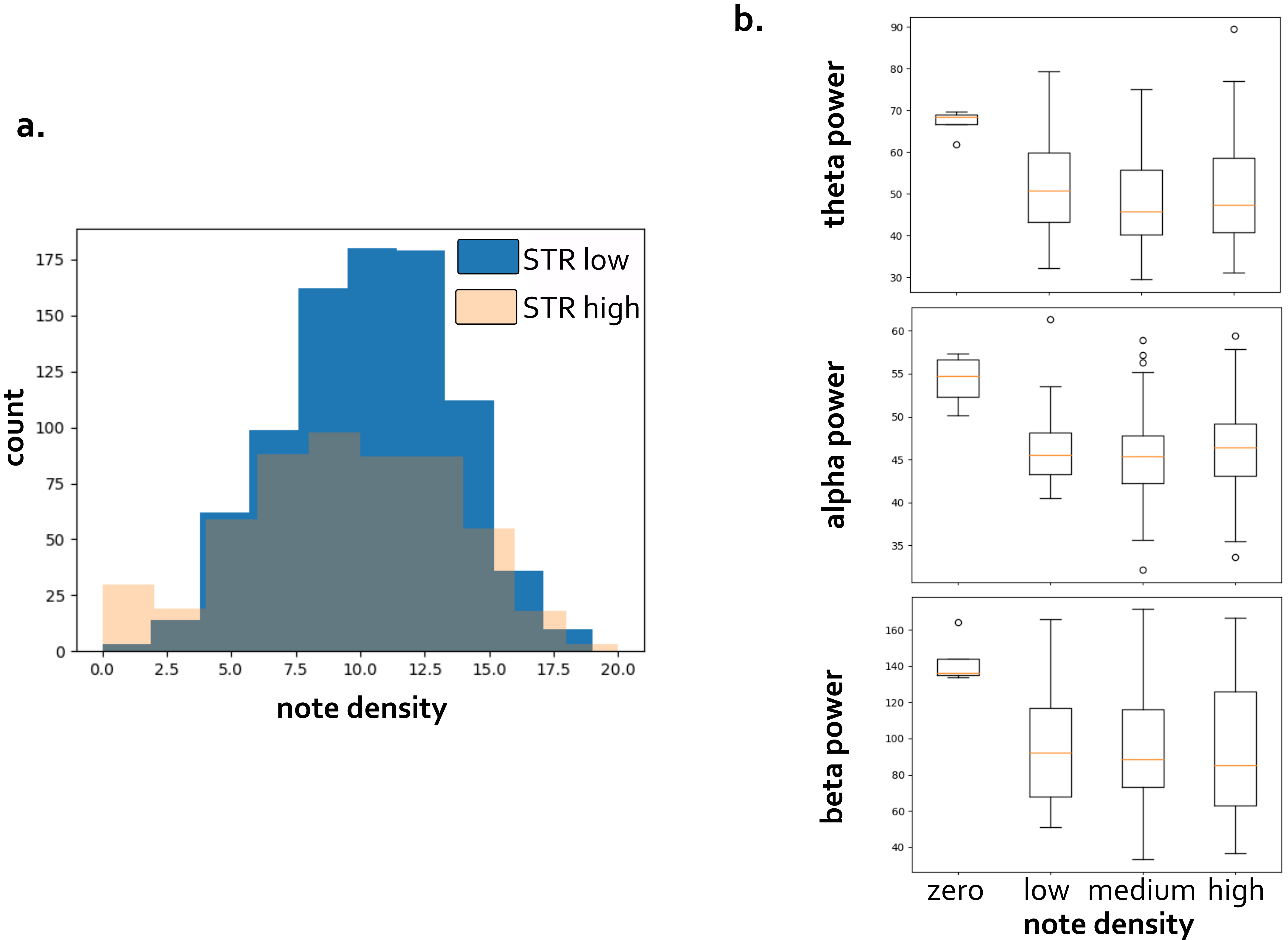
